# Faciotopy—a face-feature map with face-like topology in the human occipital face area

**DOI:** 10.1101/012989

**Authors:** Linda Henriksson, Marieke Mur, Nikolaus Kriegeskorte

## Abstract

The occipital face area (OFA) and fusiform face area (FFA) are brain regions thought to be specialized for face perception. However, their intrinsic functional organization and status as cortical areas with well-defined boundaries remains unclear. Here we test these regions for “faciotopy”, a particular hypothesis about their intrinsic functional organisation. A faciotopic area would contain a face-feature map on the cortical surface, where cortical patches represent face features and neighbouring patches represent features that are physically neighbouring in a face. The faciotopy hypothesis is motivated by the idea that face regions might develop from a retinotopic protomap and acquire their selectivity for face features through natural visual experience. Faces have a prototypical configuration of features, are usually perceived in a canonical upright orientation, and are frequently fixated in particular locations. To test the faciotopy hypothesis, we presented images of isolated face features at fixation to subjects during functional magnetic resonance imaging. The responses in V1 were best explained by low-level image properties of the stimuli. OFA, and to a lesser degree FFA, showed evidence for faciotopic organization. When a single patch of cortex was estimated for each face feature, the cortical distances between the feature patches reflected the physical distance between the features in a face. Faciotopy would be the first example, to our knowledge, of a cortical map reflecting the topology, not of a part of the organism itself (its retina in retinotopy, its body in somatotopy), but of an external object of particular perceptual significance.

## 1. INTRODUCTION

The human ventral stream contains macroscopic regions that respond selectively to certain categories, including faces and places (Epstein & Kanwisher, 1998; Kanwisher, McDermott, & Chun, 1997). Domain-specific computational mechanisms (Kanwisher, 2000) might be required to meet the difficult computational challenge of visual object recognition. The particular category preferences found appear broadly consistent with the behavioural importance of the ability to recognize faces and places. However, we do not yet understand the computations performed in these category-selective regions or their intrinsic spatial organisation. In addition, a longstanding debate has yet to be resolved about the question whether these regions form visual areas (Felleman & Van Essen, 1991; Van Essen criteria for visual areas) and functional modules (Kanwisher, 2000) or merely peaks of selectivity within a single more comprehensive object-form topography (Haxby et al., 2001).

A prominent theory of the global layout of the ventral stream states that regions selective for faces and places start out in development as a retinotopic protomap (Hasson, Harel, Levy, & Malach, 2003; Hasson, Levy, Behrmann, Hendler, & Malach, 2002; Levy, Hasson, Avidan, Hendler, & Malach, 2001). Through experience, each patch of cortex develops selectivity for the visual shapes that most often appear in the retinal region it represents. Faces often appear at the fovea, because we tend to fixate them and because their retinal size is only a few degrees visual angle when viewed at typical distances. The central part of the retinotopic protomap, according to the theory, therefore turns into the fusiform face area (FFA; Kanwisher et al., 1997; Puce, Allison, Asgari, Gore, & McCarthy, 1996) and the occipital face area (OFA; Gauthier et al., 2000). Places and scenes, by contrast, are more physically extended and typically occupy a wide visual angle. The parahippocampal place area (Aguirre, Detre, Alsop, & D’Esposito, 1996; Epstein & Kanwisher, 1998) therefore develops in the peripheral part of the protomap, according to the theory.

Here we investigate the hypothesis that the same principle explains the intrinsic spatial organisation of face-selective regions. Let’s start with an oversimplification and imagine faces always appeared frontally at the same distance and in the same retinal location (*e.g.*, fixated centrally). A retinotopic protomap whose receptive fields cover face parts would then be expected to acquire selectivities corresponding to face parts, including the eyes, nose, and mouth. Moreover, the spatial organization of the parts would resemble the spatial layout of a face, with the nose represented in a cortical patch that lies somewhere in between the patches representing the eyes and the mouth. We refer to this kind of cortical map of face-feature detectors as faciotopic. Whereas a retinotopic map is a cortical representation whose topology resembles that of the retina, a faciotopic map is a cortical representation whose topology resembles that of a face.

The scenario sketched above was an oversimplification. In natural experience, faces are viewed at a variety of distances, and they are not always fixated centrally. In order to test whether the faciotopy hypothesis is even plausible when we consider more natural viewing conditions, we used a simple simulation (Figure 1). For each face feature, we estimated the spatial distribution of retinal exposures when viewing conditions were drawn randomly from realistic distributions of viewing distances and fixation points. This gave us the spatial distribution on the retina of mouth exposures, for example, and a similar distribution for each other face feature. Despite the variability in viewing conditions, the peaks of the retinal feature exposure maps still formed a map of a face. This suggests that a retinotopic protomap with a receptive field size roughly corresponding to face parts might develop into a faciotopic map if its patches acquire selectivity for the features they are most frequently exposed to—even when viewing conditions are quite variable. Although our simulation did not include variations of viewing angle, it models a substantial part of our visual experience with faces and convinced us that the faciotopy hypothesis is plausible and merits an empirical investigation with functional magnetic resonance imaging (fMRI).

**Figure 1.**
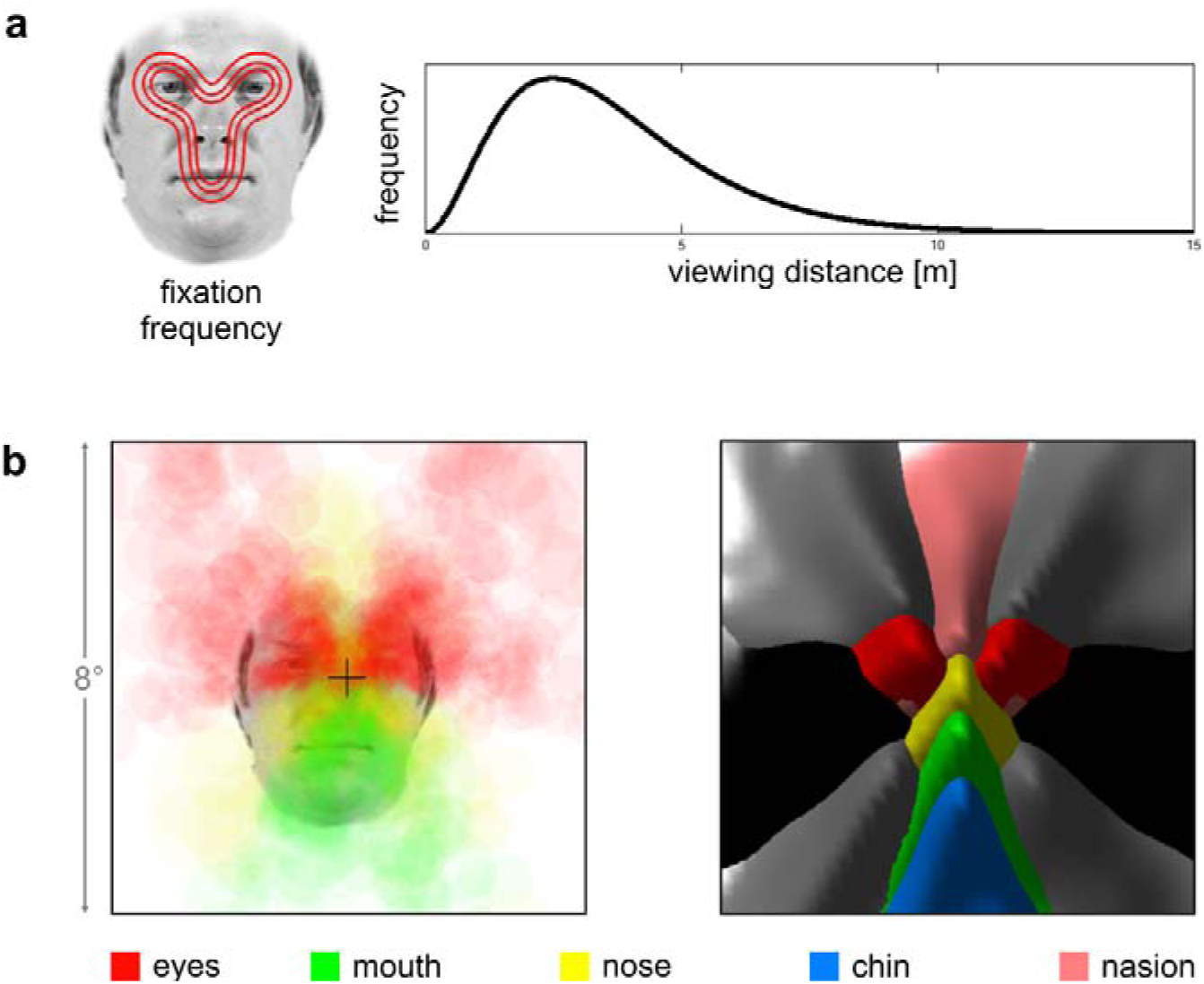
Toy simulation of retinal feature exposure in natural experience. A simple simulation suggests that although each face feature will fall in a wide range of retinal positions in natural experience, retinal feature exposure maps still reflect the geometry of a face. a) The left panel shows an idealized frequency distribution of fixation locations on faces. The distribution is based on face fixation measurements from the literature (Hsiao & Cottrell, 2008; van Belle, Ramon, Lefevre, & Rossion, 2010) and visualized by three isoprobability-density contours (red) on an example face. The right panel shows an assumed distribution of viewing distances (Gamma distribution). b) We simulated natural retinal exposure to faces as random independent draws from the fixation-location distribution and the viewing-distance distribution, assuming that the height of a face is 12.5 cm (chin to eye brows). Each draw from the distribution exposes the retina (and thus the cortical retinotopic protomap) to each of the face features at a certain location. Both panels show retinal maps of natural face feature exposure. In the left panel, the feature exposures are visualized on the retinal map (fovea indicated by cross) by transparent disks whose size reflects the size of the face (resulting from the viewing distance) and whose color codes the face feature (color legend at the bottom). In the right panel, the size of the retinal face projection is ignored and the exposure frequency distribution over the retina is visualized for each feature (colors) as a surface plot (frequency axis pointing out from picture plane). The right panel includes all 12 features used in this study. Gray and black code for the outer face features: hairlines, ears, and lower cheeks, as shown in Fig. 2a.

The FFA and OFA respond to faces even when they are presented peripherally (Hasson et al., 2003). Even if selectivities to certain types of natural shapes develop from a retinotopic protomap, the resulting shape detectors might acquire substantial tolerance to retinal position. We hypothesized that within a faciotopic map, similarly, each feature detector will respond to its preferred feature with some level of tolerance to the precise retinal position. In this study, we tested the faciotopy hypothesis by presenting images of isolated face features to subjects during fMRI scanning. To avoid confounding faciotopy with retinotopy, all face features were presented centrally at fixation. Results suggest that OFA, and to a lesser extent also FFA, is organized into a faciotopic map.

## 2. MATERIALS AND METHODS

### 2.1 Subjects

Thirteen healthy volunteers (6 females, age range 20–45) with normal or corrected-to-normal vision took part in this study. Data from one subject had to be excluded from the analysis due to technical difficulties during data acquisition (scanner failure). Ethical approval for the research was obtained from Cambridge Psychology Research Ethics Committee (CPREC). Subjects gave written informed consent before participating in the study.

### 2.1 Face-feature stimuli and experimental design

The stimuli were images of face features which had been sampled from high-resolution frontal face photographs of 92 individuals. The faces in the photographs were first aligned using Matlab by manually marking the midpoints of the eyes and the mouth in each image, then finding the rigid spatial transformation between these points and applying the transformations to the images. From the aligned face images, the following twelve face features were sampled using equal-sized, non-overlapping windows (Fig. 2a): left and right eye, the space between the eyes, nose, mouth, left and right hairline, left and right ear, left and right jaw line, and chin. The vertical positions of the sampling windows for the ears needed to be manually adjusted to match the individual variability in the position of the ears but all other features were sampled using the same windows for each face.

**Figure 2.**
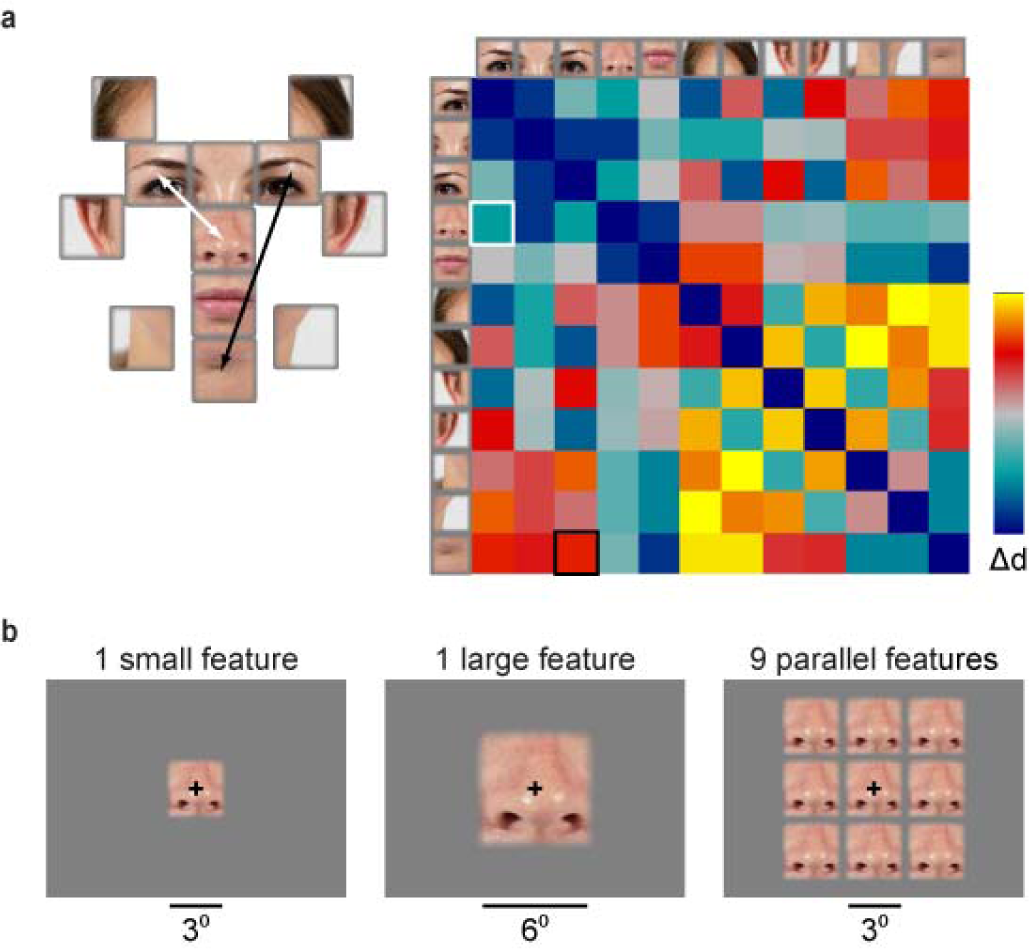
Face-feature stimuli. a) Twelve face features were sampled from 92 frontal face photographs. The sampling windows were equal-sized and non-overlapping. The elements in the matrix reflect the relative physical distances between the features. b) All face features were shown at the centre of the visual field. Three different stimulus layouts were used: a single small feature, a single large feature, and nine small features presented in parallel.

We used three different spatial layouts for stimulus presentation (Fig. 2b): one small feature (image diameter: 3°) presented at the centre of the screen, one large feature (image diameter: 6°) presented at the centre of the screen, and nine small features (all the same) presented in parallel. The subjects fixated a black cross at the centre of the screen throughout the experiment. The stimuli were shown in a blocked fMRI design, where during one 16-second stimulation-block, 16 different exemplars of the same face feature were presented (*e.g.*, 16 noses sampled from different individual faces). Each image was shown for 750 ms with a 250 ms fixation baseline between the different feature exemplars. Each experimental run consisted of two blocks for each of the face features, and every fifth block was a baseline block with the fixation cross presented alone. The total duration of an experimental run was approximately 9 minutes. Subjects attended two measurement sessions with two experimental runs for each stimulus layout within each session. The presentation order of the face features within each run and the order of the spatial-layout runs within a session were pseudorandomized and balanced across the subjects and between the two measurement sessions for each subject. The stimuli were created with Matlab, and their timing was controlled with Presentation (Neurobehavioral Systems). The stimuli were projected with a Christie video projector to a semitransparent screen, which the subjects viewed via a mirror. The subjects were familiarized to the stimuli and task before the experiments.

To direct subjects’ attention to the stimuli during the experiment, they performed a task on the stimuli. The sampling window of the face feature was displaced by half of its width 1–3 times within a stimulus block (*e.g.*, nose not shown at the centre of the visual field but shifted to the left from the fixation cross; the position and size of the stimulus images remained the same however), and the subjects were instructed to press a button when detecting these displacements of the features.

### 2.3 Regions-of-interest

The primary visual cortex (V1) was localized in each individual based on the cortical folds via a surface-based atlas alignment approach developed by Hinds et al. (2008). Peripheral V1 was excluded from the ROIs based on the spatial extent of the overall fMRI response to the face-feature stimuli. Occipital face area (OFA) and fusiform face area (FFA) were localized based on independent functional localizer data. During the functional localizer run, the subjects were presented with blocks of images of faces (different from the faces used for sampling the face features), scenes, objects, and phase-randomized textures. Subjects performed a one-back task on the stimulus images.

### 12.4 Data acquisition and analysis

Functional and anatomical MRI data were acquired using a 3T Siemens Tim Trio MRI scanner equipped with a 32-channel head coil. During each main experimental run, 252 functional volumes were acquired using an EPI sequence with imaging parameters: repetition time 2.18 s, 35 slices with 2 mm slice thickness (no gap), field of view 192 mm × 192 mm, imaging matrix 96 × 96, echo time 30 ms, and flip angle 78°. Each subject attended two measurement sessions with six main experimental runs in each (two runs for each stimulus layout, Fig. 2b), and one functional localizer run at the end of each session. Two high-resolution structural images were acquired in the beginning of the first measurement session using an MPRAGE sequence, from which the white and gray matter borders were segmented and reconstructed using Freesurfer software package (Dale, Fischl, & Sereno, 1999; Fischl, Sereno, & Dale, 1999). One structural image was acquired in the beginning of the second measurement session to co-register the data between the two sessions.

Functional data were pre-processed with SPM8 (Wellcome Department of Imaging Neuroscience) Matlab toolbox. The first four functional images from each run were excluded from the analysis to reach stable magnetization. The functional images were corrected for interleaved acquisition order and for head motion. The data from the second measurement session were co-registred and re-sampled to the same space with the first measurement session data. For the linear discriminant analysis, the data were also spatially smoothed using a 4 mm Gaussian smoothing kernel. All analysis were performed in the native space (no normalization was applied).

We estimated the responses for the face-feature stimuli using general linear model (GLM) analysis as implemented in SPM8. The onsets and durations of the stimulus blocks were entered as regressors-of-interest to the GLM, and convolved with the canonical hemodynamic response model. Additional regressors included the timings of the task images and the six head-motion-parameters. During the parameter estimation, the data were high-pass filtered with 300-s cut-off, and serial autocorrelations were estimated with restricted maximum likelihood algorithm using a first-order autoregressive model. For representational similarity analysis, the parameter estimates were transformed into t values.

### 2.5 Linear discriminant analysis

We used linear discriminant analysis (Kriegeskorte, Formisano, Sorger, & Goebel, 2007; Nili et al., 2014) to study the discriminability of the response patterns evoked by the different face-feature stimuli. The data were first divided into two independent sets based on the measurement session. For each pair of face-feature stimuli, Fisher linear discriminant analysis was applied to find the weights for the voxels that discriminated between the response patterns and then the weights were applied to the independent data to calculate the linear-disciminant t-value, reflecting the discriminability between the response patterns evoked by two different face-features. The analyses were done on individual data, and the linear-discriminant t-values were pooled across the twelve subjects and converted to p-values. All pairwise comparisons of the face-features were collected to matrices; multiple testing (66 pairwise comparisons of 12 face features) was accounted for by controlling the false-discovery rate.

To test for size-tolerance of the face-feature representations, the Fisher linear discriminant was fit to the response patterns evoked by the small face-feature images and tested on the response patterns evoked by the large face-feature images.

### 2.6 Representational similarity analysis

To characterize the face-feature representations in each ROI, we computed the dissimilarities between the response patterns evoked by the face-feature stimuli and compared them with model predictions of the representational distances (Kriegeskorte, Mur, & Bandettini, 2008; Nili et al., 2014). For each ROI, the dissimilarities between the response patterns were assembled in a representational dissimilarity matrix (RDM; a brain RDM), where each value reflects the representational distance between two face-feature stimuli. Our measure of response-pattern dissimilarity was correlation distance (1 - Pearson linear correlation). For each individual, the RDMs were calculated separately from the response patterns for each stimulus layout (Fig. 2b) and for the two measurement sessions. For the comparison of the brain representation to the model representations, the RDMs for the small and large face-features from the two measurement sessions were averaged.

The face-feature representations in V1, OFA and FFA were compared to three predictions of the representational distances between the face features: 1) Gabor wavelet pyramid (GWP) model, 2) physical distances between the face features in a face (physical distance reference, Fig. 2a), and 3) physical distances between the face features when symmetric face features are represented in same locations (symmetric reference). The GWP model captures the low-level image similarities between the face-feature stimuli, and was adopted from Kay et al. (2008). Figure 2a shows the physical distance reference matrix, where the values are the distances between the face-feature sampling windows. The distance matrix captures the spatial relationships between the features in a face. The symmetric reference was otherwise identical to the matrix shown in Figure 2a, but the distances between the symmetric features (eyes, ears, hairlines, jaw lines) were 0 and the distances from the two symmetric features to the other features were the same.

**Figure 3.**
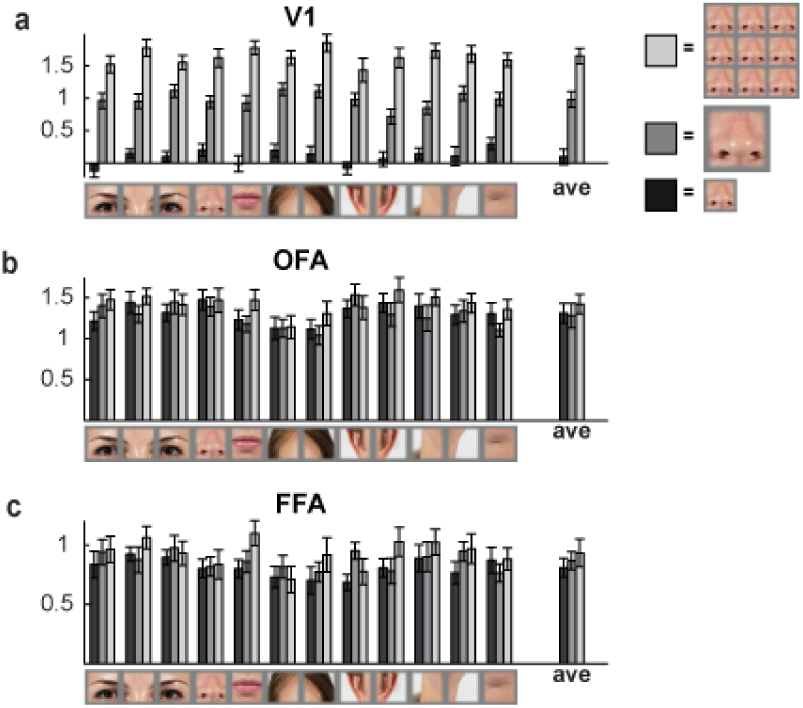
Mean responses to different face-feature stimuli. Mean responses for the 12 different face-feature stimuli are shown separately for the three different conditions (black = 1 small feature, gray = 1 large feature, light gray = 9 parallel features) in (a) V1, (b) OFA, and (c) FFA. The error-bars indicate SEMs across the 12 subjects.

We tested the relatedness between the model and brain RDMs by comparing the rank orders of the dissimilarities using Kendall’s tau-a rank correlation (for details, please see Nili et al., 2014). The relatedness of each of the model RDMs (GWP, physical distance reference, symmetric reference) to a brain RDM was tested using one-sided signed-rank test across the single subject RDM correlations. To evaluate differences between the relatedness of the model RDMs to a brain RDM, the difference between the RDM correlations of two models in each subject was calculated and tested using a two-sided signed-rank test across the subjects. This was repeated for each pair of models and the multiple testing was accounted for by controlling the false-discovery rate. A noise ceiling of the expected RDM correlation was estimated for each brain region as described by Nili et al. (2014).

The relationships of the model and brain RDMs were visualized using multidimensional scaling (Nili et al., 2014). The first step is to build a matrix of the pairwise correlations (1 - Kendall’s tau-a rank correlation) between all brain and model RDMs. To avoid the contribution of intrinsic fluctuations inflating the representational similarity between two brain regions (Henriksson, Khaligh-Razavi, Kay, & Kriegeskorte, 2015), the RDMs of the visual areas were compared between RDMs constructed from response patterns from different measurement sessions. The multidimensional scaling arrangement of the (dis)similarity matrix of the RDMs provided a visualization of the relatedness of the face-feature representations in different visual areas, and between the visual areas and models.

### 2.7 Face feature map estimation

Finally, we tested whether the face-feature representations reflect faciotopy, that is, whether the cortical distances between the representations of different face features were explainable by the physical distances between the features in a face. Within each ROI (left and right V1, left and right OFA, left and right FFA), we estimated a single location for each face feature using the following approach. For each voxel, we determined which feature was preferred (highest t value) over the other features at that voxel. We then considered the local spherical neighbourhood around each voxel in an ROI like OFA, and assigned the voxel the feature that was most frequently the preferred feature in the neighbourhood. We then looked for the voxel with the highest feature preference (defined as the number of times the feature was preferred in the local neighbourhood) and assigned that voxel together with its local neighbourhood to that feature. This procedure was repeated until all features had a cluster of voxels or all above-threshold voxels had been assigned to features. The size of the neighborhood (radius of a sphere) and the T-value threshold were optimized by evaluating the replicability of the distance matrix across the two measurement sessions (no assumption of faciotopy, only for replicable distance matrix between the face-feature locations). The feature-preference clusters were searched for in 3D space (voxel coordinates) and assigned to the cortical surface of the individual. All pair-wise distances between the cortical patches assigned to the face features were calculated along the cortical surface and assembled in a matrix similar to the distance matrix shown in Fig. 2a. To test for faciotopic representation, the patch-distance matrix was compared to the matrix of the physical distances between the features in a face. This analysis was identical to the representational similarity analysis of the response-pattern dissimilarity matrices and model RDMs.

## 3. RESULTS

### 3.1 V1, OFA, and FFA respond to isolated face features

We measured fMRI responses to visual presentations of 12 isolated face-features (Fig. 2a): left and right eye, the space between the eyes, nose, mouth, left and right hairline, left and right ear, left and right jaw line, and chin. The features were extracted from frontal face photographs using equal-sized, non-overlapping sampling and were presented in a blocked fMRI design with three conditions (Fig 2b): one small feature shown at the centre of the screen, one large feature shown at the centre of the screen, and nine small features presented in parallel. The small stimulus (3° diameter) was selected to roughly correspond to the size of the face features at a viewing distance of 1 m, whereas the simultaneous presentation of nine small features could be an optimal stimulus for a feature detector. Two face-selective regions-of-interest (OFA, FFA) and the primary visual cortex (V1) were defined in each hemisphere in each subject based on independent localizer data. The results from the left and right hemisphere were averaged.

Figure 3 shows the mean fMRI response strengths for the face-feature stimuli in V1, OFA and FFA. The different conditions (small features, large features, 9 parallel features) are shown in different shades of gray. The V1-ROI covered eccentricities up-to the size of the 9-parallel-features stimulus, and thus it is expected that in V1 the small stimulus evokes the smallest response and the largest stimulus (9 parallel features) evokes the largest mean response (Fig 3a). More interestingly, in OFA and FFA, this retinotopic effect was largely abolished and the three sizes of the face-feature stimuli evoked approximately equal-sized responses. The only exception is the mouth stimulus that evoked a larger response both in OFA (p = 0.027; signed-rank tests) and in FFA (p = 0.016; signed-rank tests) when presented in the nine parallel feature configuration compared to the one small feature presented at the centre of the screen. Overall, each face-feature stimulus evoked a clear response in all regions-of-interest.

### 3.2 V1, OFA, and FFA response-patterns distinguish the face features

We have now shown that V1, OFA and FFA respond to isolated face-features (Fig. 3), but do they also discriminate between the face features (*e.g.*, an eye from a mouth)? Figure 4a shows the results from linear discriminant analysis (Nili et al., 2014): the discriminability of each pair of face-feature stimuli was evaluated by fitting a Fisher linear discriminant to the response patterns from the first fMRI session and by testing the performance on the response patterns from the second fMRI session (same subject, different day, different stimulus presentation order, all stimulus layouts). The analyses were done on individual data and the results were pooled across the twelve subjects. The left column in Figure 4a shows the linear-discriminant t-values, reflecting the discrimability of each pair of face-feature stimuli from the response patterns, and the right column shows the corresponding p-values. In V1, the response patterns discriminated each pair of face-feature stimuli, except the two hairlines from each other and the mouth from the chin (Fig. 4a, first row). In OFA, each pair of the face-feature stimuli could be discriminated from the response patterns (Fig. 4a, middle row). In addition, there appears to be a distinction between the inner (first five elements in the linear discriminant t-value and p-value matrices; *e.g.*, the eyes) and outer face-features (elements 6–12 in the matrices; *e.g.*, the ears), that is, the t-values are high for the discriminability of these stimulus pairs in OFA, and also in FFA. Moreover, in FFA, the symmetric face-features (the eyes, the hairlines, the ears, the jaw lines) evoked indistinguishable response patterns (bottom row in Fig. 4a; see the blue rectangles in the p-value matrix).

**Figure 4.**
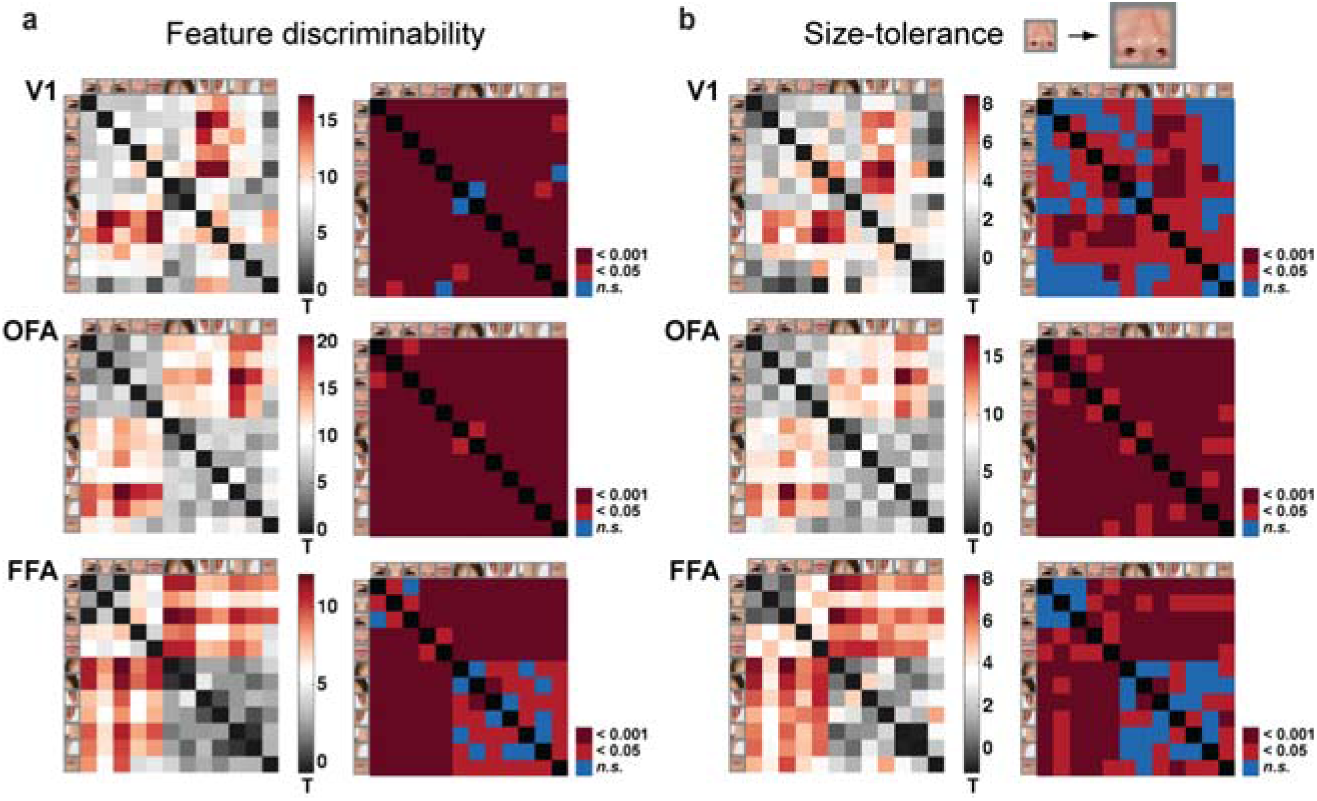
Distinctness and size-tolerance of the face-feature representations in V1, OFA and FFA. a) The linear discriminant analysis t-values and the corresponding corrected p-values are shown for all pair-wise comparisons of the face-feature response-patterns in V1 (top panel), OFA (middle panel) and FFA (bottom panel). The training data was the data from the first measurement session and the testing was done on data from the second session. In OFA, all face-feature stimuli could be discriminated from each other (no blue squares in the p-value matrix). b) To test for size-tolerance of the feature representations, the linear discriminant analysis was performed by training the classifiers on the small face-feature response-patterns and testing the classifiers on the large face-feature response-patterns. The results are shown as in (a). OFA shows successful generalization of the face-feature discrimination across stimulus size (no blue squares in the p-value matrix), suggesting size-tolerant feature representations.

### 3.3 OFA discriminates every pair of face features with tolerance to the feature size

The use of both small and large features as stimuli enabled us to study the size-tolerance of the face-feature representations in V1, OFA and FFA. In general, a true higher-level representation of an object category should show tolerance to identity-preserving image transformations, such as scaling the image size. Figure 4b shows the results from Fisher linear discriminant analysis when the classifier was trained to distinguish the response patterns for the small face-feature stimuli and the testing was done on the response patterns for the large stimuli—successful decoding would imply generalization across stimulus size and hence size-tolerance. Most importantly, OFA response-patterns discriminated between each pair of face-feature stimuli (middle row in Fig. 4b), indicating size-tolerant face-feature representations in OFA. In V1, the classifier’s performance was much worse than with the same-sized stimulus images (cf. top rows in Figs. 4a and 4b). The performance in FFA was also impaired by the use of different sized stimulus images for training and testing. In FFA, however, the distinction between the inner and outer face features was preserved. Results on the generalization from the small features to the nine parallel features, and from the large features to the nine parallel features are shown in Supplementary Figure 1.

### 3.4 OFA response-pattern dissimilarity structure is better explained by the physical distance between the face features than low-level image properties, whereas the opposite is true for V1

Next we characterized the face-feature representations in V1, OFA and FFA using representational dissimilarity matrices (RDMs; Kriegeskorte et al., 2008; Nili et al., 2014), which compare the response patterns elicited by the stimuli—here each value in an RDM reflects the representational distance between two face-feature stimuli. RDMs can be directly compared between two brain regions by computing the rank-correlation between their RDMs; if two brain regions represent the stimuli identically, the ordering from the most similar stimulus-pair to the least similar stimulus-pair is the same (high rank correlation). This comparison should, however, be done on independent trials to avoid the contribution of intrinsic cortical dynamics inflating the representational similarity between brain regions (Henriksson et al., 2015). Moreover, a brain RDM can directly be compared to an RDM constructed based on model predictions of the similarity of the representations between the stimuli.

We compared the brain RDMs to three models: physical distances between the face features in a face (physical distance reference; Fig. 2a), physical distances between the face features when symmetric face-features are represented in the same locations (symmetric reference), and Gabor wavelet pyramid (GWP) model. Figure 2a shows the physical distance reference matrix, where each value in the matrix reflects the distance between two features in a face. The symmetric reference matrix was otherwise identical to the matrix shown in Figure 2a, but the distance between the symmetric face features was zero and the distances from two symmetric face features to other features was identical. The GWP-model captures the low-level image properties of the stimuli (edges, for example, at the same locations in the stimulus images would be predicted to elicit a similar response in low-level visual areas).

Figure 5a shows the results from the comparison of the brain RDMs to the three model RDMs. The brain RDMs were constructed from the response patterns to the small and large stimulus images (see Methods for details); the results are shown separately for the different stimulus layouts in Supplementary Figure 2. The V1 RDM of the face-feature stimuli was best explained by the GWP model (p < 0.001; one-sided signed-rank test), reflecting the low-level image properties of the stimulus images. The GWP model explained the V1 representation better than the face-feature physical distance matrix or the symmetric distance matrix (FDR q<=0.05; two-sided signed-rank tests; FDR-corrected for multiple comparisons). In OFA, both the face-feature physical distance matrix and the symmetric distance matrix explained variance in the representation (p < 0.01; one-sided signed-rank tests). In addition, the physical distances between the face-features explained the representation better than the low-level image properties captured by the GWP model (FDR q<=0.05; two-sided signed-rank test; FDR-corrected for multiple comparisons). A similar trend was observed in FFA, where the physical distance matrix and the symmetric distance matrix both explained variance in the representation.

**Figure 5.**
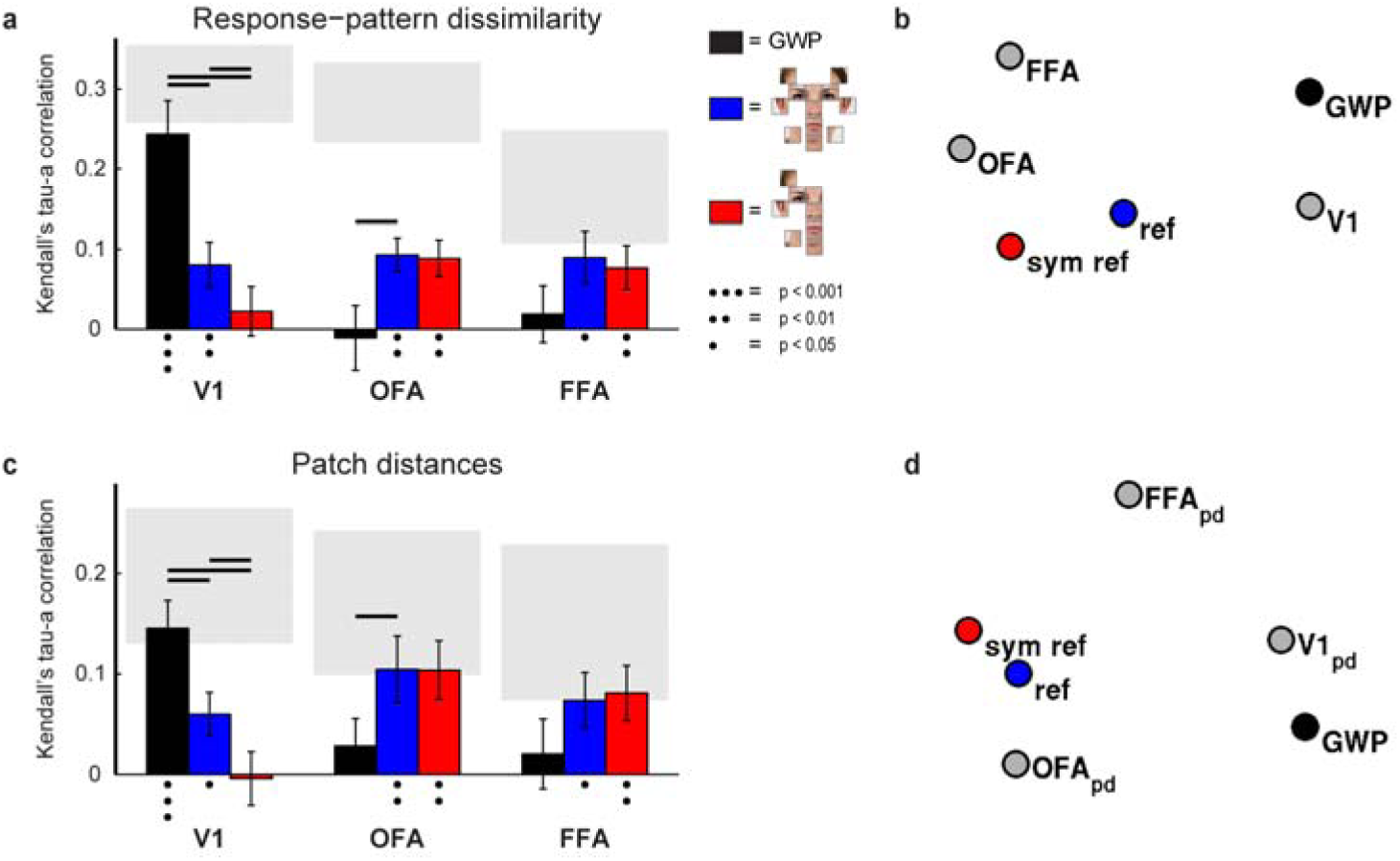
Evidence for faciotopic representation in OFA. a) The face-feature representations in V1, OFA and FFA, as reflected in the response-pattern dissimilarities, were compared to three models: Gabor wavelet pyramid model (GWP; black bars) of the low-level image similarity between the face-feature images, physical distances between the face-features in a face (blue bars), and physical distances between the face-features when symmetric face-features have a single, overlapping representation (red bars). The gray rectangles are estimates of noise ceiling (Nili et al., 2014). The error-bars indicate SEMs across the 12 subjects. The dots below the bars indicate that the model significantly explained variance in the brain representation (one-sided signed-rank test across the single-subject RDM correlations). Significant difference between the models’ relatedness to the brain representations are indicated with the black lines (two-sided signed-rank test across subjects, multiple testing accounted for by controlling the false discovery rate at 0.05). Supplementary Figure 3 shows the results separately for the left and right hemispheres. b) Multidimensional-scaling visualization of the relationships between the brain representations (as reflected in the response-pattern dissimilarities) and the three models is shown (dissimilarity: 1 – Kendall’s tau-a rank correlation, criterion: stress). c) A single patch of cortex was estimated for each face-feature based on the feature preferences. The relative distances between the face-feature-preference patches were compared to the three model representations. In OFA, the physical distances between the face-features significantly explain the distances between the face-feature-preference patches along the cortical surface (blue bars), reaching the noise ceiling of the patch-distance estimates. d) Multidimensional-scaling visualization of the relationships between the estimated face-feature patch-distances (pd), and the three models is shown (dissimilarity: 1 – Kendall’s tau-a rank correlation, criterion: stress).

Figure 5b shows a multidimensional-scaling visualization of the relationships between the three models and the V1, OFA and FFA representations, as captured by the response-pattern dissimilarity matrices. The distances reflect the correlation distance between the RDMs; that is, how similar the representations are. The OFA and FFA representations are more similar to each other than to the V1 representation. The V1 representation was most similar to the GWP model whereas the OFA representation was more similar to the face-feature physical distance models.

### 3.5 Distances between face-feature-preference patches suggest faciotopy in OFA

Thus far we have shown that especially in OFA the response-pattern dissimilarities do reflect the physical distances between the face features (Fig. 5 a–b). For the underlying representation to be truly faciotopic, the distances between cortical locations with preference for a specific feature would also reflect the topology of the face features in a face. To test for a faciotopic representation, we estimated for each face feature a single location on the cortex within each ROI and calculated the distances between these face-feature preference patches along the cortical surface. All pair-wise distances between the face-feature preference patches were collected to a matrix similar to the reference matrix shown in Figure 2a.

Figure 5c shows the results how well the distances between the face-feature preference patches along the cortical surface were explained by the physical distances between the face-features in a face (blue bars) or by the symmetric map where symmetric face-features have overlapping representations (red bars). The results are consistent with the response-pattern dissimilarity results shown in Figure 5a. In V1, the distances between the “face-feature patches” were better explained by the low-level image properties between the stimulus images as captured by the GWP model than by the physical distances between the face-features (FDR q<=0.05, two-tailed signed-rank test; FDR-corrected for multiple comparisons). The opposite was true for OFA, where the physical distances between the face features best explained the distances between the face-feature preference patches (FDR q<=0.05, two-tailed signed-rank test; FDR-corrected for multiple comparisons). Figure 5d shows a multidimensional-scaling visualization of the relationships, where the distances reflect the similarity of the representations. The physical distances between the face features in a face best explain the OFA representation, as reflected in the cortical distances between the face-feature-preference patches, suggesting faciotopy in OFA.

## 4. DISCUSSION

Our hypothesis was that face-selective regions in human ventral cortex might be organized into faciotopic maps, in which face feature detectors form a map whose topology matches that of a face. Faces, and especially the eye region, are frequently fixated from an early age (Farroni, Csibra, Simion, & Johnson, 2002; Goren, Sarty, & Wu, 1975), and a retinotopic protomap (Hasson et al., 2003; Levy et al., 2001) could develop into a faciotopic map if patches acquired selectivity for the face features that they are most frequently exposed to. We first performed a simple simulation to support the faciotopy hypothesis, and then measured fMRI responses to isolated face features, all presented foveally to prevent the results from being driven by retinotopy. First, we reported that V1, OFA, and FFA respond to isolated face features and their response patterns distinguish the different face-feature stimuli. Both OFA and FFA emphasized the distinction between the inner (*e.g.*, eyes, nose, mouth) and outer face features (*e.g.*, ears, chin, hairline) in their representations. Furthermore, the face-feature discrimination was tolerant to a change of feature size in OFA. Size tolerance was smaller in FFA and absent in V1. We then tested for each region how well the observed response-pattern dissimilarities could be explained by each of three models: a physical-distance model (Fig. 2a), a mirror-symmetric physical-distance model, and a Gabor wavelet pyramid model. The first two models reflect natural face topology; the latter captures the low-level image properties of the stimuli. Our results indicate that the response-pattern dissimilarity structure in OFA is better explained by the physical distances between the face features than by low-level image properties, whereas the opposite is true for V1. Results for FFA were similar to OFA, but the difference between the models was weaker. Finally, a true faciotopic organization requires more than a match between pattern dissimilarities and physical face-feature distances: it requires that the map of cortical locations which preferentially respond to each face feature reflects the topology of a face. To test this, we computed cortical distances between feature-preference locations, and compared them to our three models. The distances between the cortical feature-preference patches in OFA were indeed best explained by the physical distances between the features in a face, supporting the existence of a faciotopic map in OFA.

### 4.1 Faciotopy is consistent with previous findings on OFA

The function of OFA has also previously been associated with processing of face features (e.g., Haxby, Hoffman, & Gobbini, 2000; Liu, Harris, & Kanwisher, 2010; Pitcher, Walsh, Yovel, & Duchaine, 2007). Previous studies have shown that transcranial magnetic stimulation (TMS) at right OFA disrupts face-feature discrimination (Pitcher et al., 2007) and that OFA is activated more by a face with real inner face-features present than a face with the features replaced by black ovals (Liu et al., 2010). Moreover, the face-selective neurons in the macaque posterior lateral face patch (PL), the likely homologue of human OFA, are driven by a single eye, especially when presented in the contralateral upper visual hemifield (Issa & DiCarlo, 2012). In the present study, we did not find any special role for the eyes over the other face features. This could be explained by the temporal resolution of fMRI; Issa et al. (2012) reported the eye-preference mainly for the early response (60–100 ms), and our fMRI responses reflect a mixture of early and late responses. In a later time-window (>100 ms), the macaque PL neurons also respond to other face features (Issa & DiCarlo, 2012).

Previous research suggests that OFA is less sensitive than FFA to the correct configuration of the face features within a face (Liu et al., 2010; Pitcher et al., 2007). This would be consistent with OFA containing a map of somewhat independently operating face-feature detectors. Results from monkey electrophysiology support a functional distinction between the two regions: the macaque posterior face patch (the putative homologue of OFA) seems to linearly integrate features (whole = sum of the parts), at least in the early response phase (Issa & DiCarlo, 2012), while the majority of neurons in the middle macaque face patch (the putative homologue of FFA) shows interactions between the face features (Freiwald, Tsao, & Livingstone, 2009). This is consistent also with a human magnetoencephalography (MEG) study showing that the early face-selective MEG response (peaking at a latency of 100 ms) reflects the presence of real face parts more strongly than the naturalness of their configuration, whereas the later response (170 ms) shows the opposite sensitivity (Liu, Harris, & Kanwisher, 2002).

### 4.2 Faciotopy and retinotopy might co-exist in OFA and other face regions

If faciotopy and retinotopy coexisted in OFA, we would expect face features presented in their typical locations to elicit the largest response. This would imply a preference of the region as a whole for a natural configuration of the features. We would also expect that the features elicit the strongest response when presented in the canonical upright position, as done here. The effect of inverting the face features remains, however, an open question. In addition, a faciotopic map would be expected to exhibit non-linear interactions to some degree when multiple face features are presented together. This would be analogous to early retinotopic cortex, which exhibits non-linear spatial interactions when multiple visual-field regions are stimulated at the same time (Pihlaja, Henriksson, James, & Vanni, 2008).

In visual field eccentricity maps (Hasson et al., 2002), OFA shows a preference for the central part of the visual field. In addition, human OFA has been shown to prefer contralateral stimuli (Hemond, Kanwisher, & Op de Beeck, 2007) and to support both position-invariant linear readout of category information and category-invariant linear readout of position information (Schwarzlose, Swisher, Dang, & Kanwisher, 2008). Although a more detailed retinotopic organization has not yet been demonstrated in human OFA, there is evidence for retinotopy in subregions of the macaque face patches (Rajimehr, Bilenko, Vanduffel, & Tootell, 2014). The reported preference for an eye-like feature in its natural visual-field position relative to fixation in monkey PL (Issa & DiCarlo, 2012) also suggests that retinotopy might play a role, and that the conjunction of the feature and its retinal location might determine the response.

In the present results, the stimulus layout with the nine parallel mouths was more effective than the single small mouth for both OFA and FFA. This could be explained by one of the nine mouths landing in the natural location below the fixation. However, we did not observe a similar preference for any other feature. The optimal fixation point across different face recognition tasks has been reported to be, on average, just below the eyes (Peterson & Eckstein, 2012), which could define the center of the faciotopic map. However, recent studies also report that face-fixation patterns, although stable within an individual, differ across individuals (Mehoudar, Arizpe, Baker, & Yovel, 2014; Peterson & Eckstein, 2013). If faciotopic maps arise from retinal face-feature exposure, they might similarly exhibit individual variability reflecting differences in individual’s preferred fixation locations.

Sensitivity to retinal position is theoretically compatible with faciotopy and expected if a faciotopic map developed from a retinotopic protomap. This is analogous to the coarser scale, where preferences for faces and places in FFA and PPA co-exist with retinotopic biases. It is all the more striking, then, that the face-feature map can be driven by centrally-presented face features independent of retinotopy. This suggests that, despite residual retinotopic biases, the face feature detectors respond with some level of position tolerance. Future studies should investigate how retinotopy and faciotopy combine in OFA and other face regions. Future studies could determine the relative contributions of retinotopy and faciotopy by systematically varying the retinal position of the presented features.

### 4.3 Can topographic maps, large and small, explain the intrinsic spatial organisation of the ventral stream?

Several pieces of evidence suggest a more global topographic representation of the human body within human occipitotemporal cortex (Bernstein, Oron, Sadeh, & Yovel, 2014; Chan, Kravitz, Truong, Arizpe, & Baker, 2010; Orlov, Makin, & Zohary, 2010; Song, Luo, Li, Xu, & Liu, 2013). Visual representations are sensitive to the configuration in which a face and a body are arranged in an image, with stronger responses for the typical configuration of face and body (Bernstein et al., 2014; Song et al., 2013) and for the typical configuration of the left and right halves of the body (right side of the body in the left visual field; Chan et al., 2010). A topographic representation of the body would be expected to devote larger areas to more informative body parts, which would include faces and perhaps also hands. The higher-level visual representations may, thus, exhibit cortical magnification of the most informative features, analogous to the early retinotopic visual areas, with their enlarged representation of the central visual field (Duncan & Boynton, 2003; Engel et al., 1994) and to the somatosensory homunculus with its enlarged representation of the parts of the organism’s own body that provide richer tactile and proprioceptive input.

Selective magnification of the most informative parts might also be a feature of a faciotopic representation. This question might be addressed in future studies that sample the face locations with higher spatial resolution in the stimulus domain and also image OFA response patterns with higher resolution, for example using high-field fMRI. The nature of the magnified representation might turn out to be quite different in body-part and face-part maps: whereas face perception is tightly coupled with social communication and ultimately relies on holistic perception of the face, visual representations of individual body parts may play an important role in tasks such as action understanding.

The spatial organisation of a cortical representation is likely to reflect not only real-world spatial regularities of the stimuli, but also functional relationships. It has been proposed that functional relationships explain the spatial organisation of motor cortex (Aflalo & Graziano, 2006; Graziano, 2006). For higher-level visual cortex, a close relationship between body-part and tool-use representations has recently been established (Bracci & Peelen, 2013).

It remains to be explored whether there are also other areas with cortical maps reflecting topology of external objects. As an object category, faces have particular perceptual significance and have prototypical configuration of features. Faces are also typically perceived in a canonical upright orientation and are fixated at particular locations. These properties make faces an ideal candidate for the development of a cortical feature map. If faciotopy develops from the central part of a retinotopic protomap (Hasson et al., 2003; Hasson et al., 2002; Levy et al., 2001), the peripheral part of the protomap might develop a topological map of the local environment in the occipital and/or parahippocampal place area. This hypothesis remains to be tested in future studies.

Overall, the idea of topographic cortical representations of face and body is consistent with an eccentricity-based protomap within the occipitotemporal cortex that develops into more specialized category representations (Hasson et al., 2002; Levy et al., 2001). Although the state of development of the cortical face-processing network in newborn infants remains to be determined, there is converging evidence that a subcortical route is responsible for the early tendency of newborns to orient towards face-like stimuli (Johnson, 2005). The innate subcortical route may be responsible for bringing faces to central visual field in newborns and might promote the development of faciotopy. Consistent with this hypothesis, visual experience during the first months after birth is necessary for normal development of configural processing of faces and, in particular, the processing of information about the spacing between the face features (Le Grand, Mondloch, Maurer, & Brent, 2001). An innate neural mechanism that triggers central fixation of faces would spatially align the repeated face exposures of a retinotopic protomap, and could explain the development of faciotopy.

## Acknowledgements

This work was supported by the Aalto University, the European Research Council (Advanced Grant #232946 to R. Hari) and the Academy of Finland Postdoctoral Researcher Grant (278957) to LH, a Netherlands Organisation for Scientific Research (NWO) Rubicon Grant to MM, and a Wellcome Trust Project Grant (WT091540MA) and a European Research Council Starting Grant (261352) to NK. The authors declare no competing financial interests.

